# Population genomic analyses of schistosome parasites highlight critical challenges facing end-game elimination efforts

**DOI:** 10.1101/2020.04.30.070656

**Authors:** Jonathan A. Shortt, Laura E. Timm, Nicole R. Hales, Zachary L. Nikolakis, Drew R. Schield, Blair W. Perry, Yang Liu, Bo Zhong, Todd A. Castoe, Elizabeth J. Carlton, David D. Pollock

## Abstract

Schistosomiasis persists in some Asian regions despite targeted end-game elimination measures. To determine the causes of this persistence, we performed reduced representation genome sequencing on *Schistosoma japonicum* miracidia collected across multiple years from transmission hotspots in Sichuan, China. We discovered strong geographic structure, suggesting that local rather than imported reservoirs are key sources of infection persistence. At the village level, parasites collected after praziquantel treatment are closely related to local pre-treatment populations. Schistosomes within villages are also highly related, suggesting that only a few parasites from a limited number of hosts drive re-infection. The close familial relationships among miracidia from different human hosts also implicates short transmission routes among humans. At the individual host level, genetic evidence indicates that multiple humans retain infections following referral for treatment. Our findings suggest that end-game schistosomiasis control measures should focus on completely extirpating local parasite reservoirs and confirming successful treatment of infected human hosts.

**One Sentence Summary:** Local reservoirs are the source for persistent schistosomiasis infections despite continuous efforts aimed at complete elimination.

## Main Text

Schistosomiasis and other human helminthiases are neglected tropical diseases that impact an estimated one billion people globally (*1-3*), causing fibrosis of the liver and bladder, anemia, and cancer (*4, 5*). China has become a model for successfully reducing schistosomiasis prevalence through aggressive control programs, such as mass administration of the anthelmintic praziquantel, that have decreased infection by 99% (*6-8*) since the 1950s. Despite this remarkable progress, transmission hotspots persist (*9-10*), with an estimated 54,000 Chinese still infected by the local species *Schistosoma japonicum* (*8*). Persistence of schistosomiasis despite control measures is also a widespread problem outside of China (*11*), underscoring the importance of addressing questions such as whether infections are acquired from local sources or imported, which hosts are sources of infections, and what is the frequency of treatment failure.

To understand transmission dynamics, explain persistence, and better inform control strategies, we analyzed genomic variation in 200 schistosome miracidia collected from 38 human hosts in 12 villages in Sichuan, China (Fig 1a). Collections occurred in 2007, 2008, 2010, and 2016, with miracidia preserved on Whatman FTA indicator cards as previously described (*12-14*). Infected human hosts were referred for praziquantel treatment following sample collection. Miracidia genomes were amplified, and ddRADseq libraries were created and sequenced as described in the literature (*14, 15*), resulting in >33,000 loci/miracidium after filtering (see Supplementary Methods, Fig S1).

**Fig. 1.**
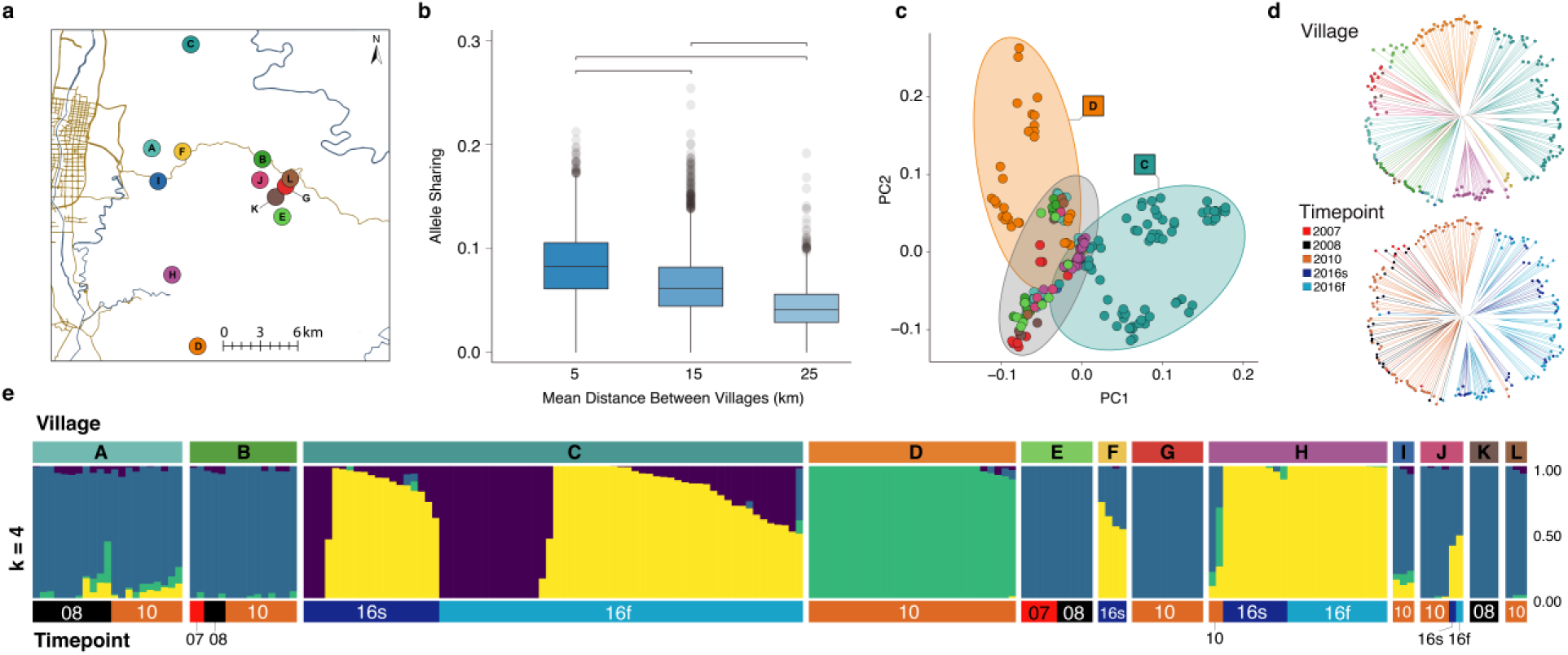
Genetic and geographic structure of schistosome miracidia sampled in Sichuan, China. a) Map showing locations of the 12 villages sampled, indicated by colored dots. b) Proportion of rare alleles shared among villages with mean, interquartile ranges, and outliers beyond the 2.5^th^ percentile shown. All comparisons were significantly different (all p < 2.2 * 10^−16^; Mann-Whitney U test). c) Principal component analysis (PCA) of genetic variation from 200 miracidia across all 12 villages. The first two principal components (PC1 and PC2) respectively account for 4.2% and 2.5% of the genetic variation among individuals. d) Neighbor-joining tree of miracidia colored by village (top) and sampling timepoint (bottom). e) ADMIXTURE plot showing optimal *k*=4 genetic clusters grouped by village and sampling timepoint. Timepoints are labeled with year of collection (e.g., 2008 in 1c or 08 in 1e); Summer and late Fall 2016 collections are labeled with small s or f, respectively.

To determine whether schistosome infections are acquired from local or regional sources, we evaluated the spatial distribution of schistosome genetic diversity across our study region. The genetic structure indicates that the parasites are more related within villages than between villages, with allele sharing decreasing significantly with geographic distance between villages (Fig 1a & 1b). Population structure is strong enough that most villages have a unique, discernible population of miracidia. For example, the first two principal components in principal components analysis (PCA) distinguish miracidia in the two most distant villages, C and D, from other villages (Fig 1c); additional principal components separate most other villages into clear clusters based on genetic similarity (Fig S2). Phylogenetic analysis of miracidia also clusters villages, with most villages occupying their own clade (Fig 1d). Estimates of population structure using *ADMIXTURE* (*16*) support this finding and identify further substructure within villages, particularly village C (Fig 1e).

The genetic structure of schistosomes within villages indicates that local infection sources were not fully eliminated by whole-village praziquantel treatments between sampling points. Miracidia from the same village fall into characteristic *ADMIXTURE* clusters regardless of sampling timepoint (Fig 1e, villages A, B, C, E, and H; see also Fig S3), and miracidia collected from the same timepoint fall into multiple clades on the phylogenetic tree (Fig 1d). However, there is a notable difference in genetic structure in village J between 2010 and 2016, the largest time span present in the data (Fig 1d and 1e). This provides clear evidence for the successful impact of control measures on population dynamics in *S. japonicum*.

To investigate infection persistence within villages, we used allele sharing to delineate familial relationships among miracidia (Fig 2a). Miracidia from the same village are closely related (Fig 1b). Posterior probabilities of relatedness were calculated from allele sharing (Supplementary Methods) and indicate that schistosome 1^st^ cousins (3^rd^ degree relatives) are extremely common within villages, but much rarer between villages (Fig 2b). As expected, pairs of miracidia collected from the same human host are often siblings (1^st^ degree relatives; Fig 2b). However, we also find a large number of 2^nd^ degree relatives within villages (Fig 2c, village D). It seems reasonable that most of these are double first cousins, given the high frequency of first cousins within villages.

**Fig. 2.**
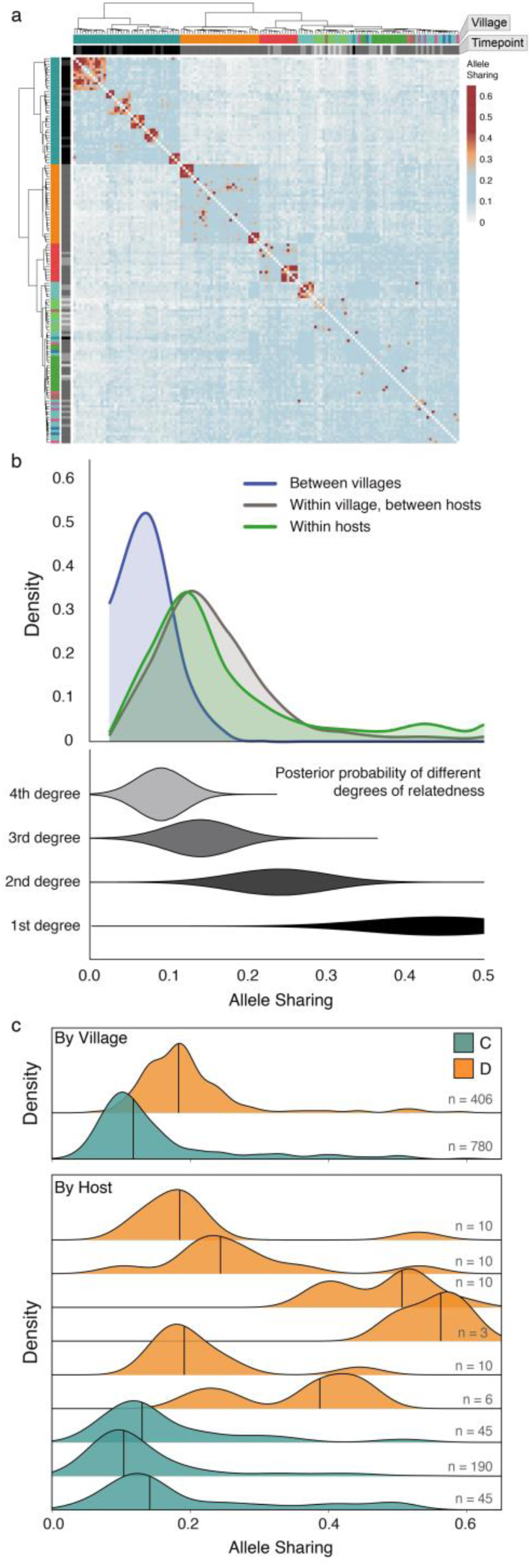
Genetic relatedness within and between villages. a) Heatmap of allele sharing between all sampled miracidia. Rows and columns are ordered using hierarchical clustering and annotated with village and timepoint. b) Distributions are shown for allele sharing between miracidia pairs sampled from different villages (blue), within villages but different hosts (grey), and within hosts (green). The posterior probabilities for different degrees of relatedness are indicated by width for 1st to 4th degree relatives in the lower plot. c) Distributions of within-village (top panel) and within-host (lower panel) allele sharing are shown for villages C (green) and D (orange). The total number of comparisons underlying each distribution is shown on the right.

High levels of allele sharing within villages indicate that parasite mate choice is often limited to relatives during the reinfection process. This limitation implies that village reinfections may have begun with a small number of infecting cercariae, likely reflecting the effectiveness of the local schistosomiasis control program (*17, 18*). However, these results suggest that long-term elimination may require identification and targeted treatment of remaining local reinfection sources.

Probabilities of relatedness among miracidia also provide detailed information on infection and transmission patterns underlying local persistence. Separate clutches of parasite siblings within individual human hosts indicate infection by multiple mating pairs, and such clutches were observed as expected (Fig 3a). We also found multiple examples of human hosts with sibling clusters that span multiple sampling time points; this is strong evidence for retained infection despite intervening treatment protocols (Fig 3b). This raises questions about the effectiveness of treatment protocols, and raises concerns that human hosts who failed to clear their infections may have served as sources of infection to others.

**Fig. 3.**
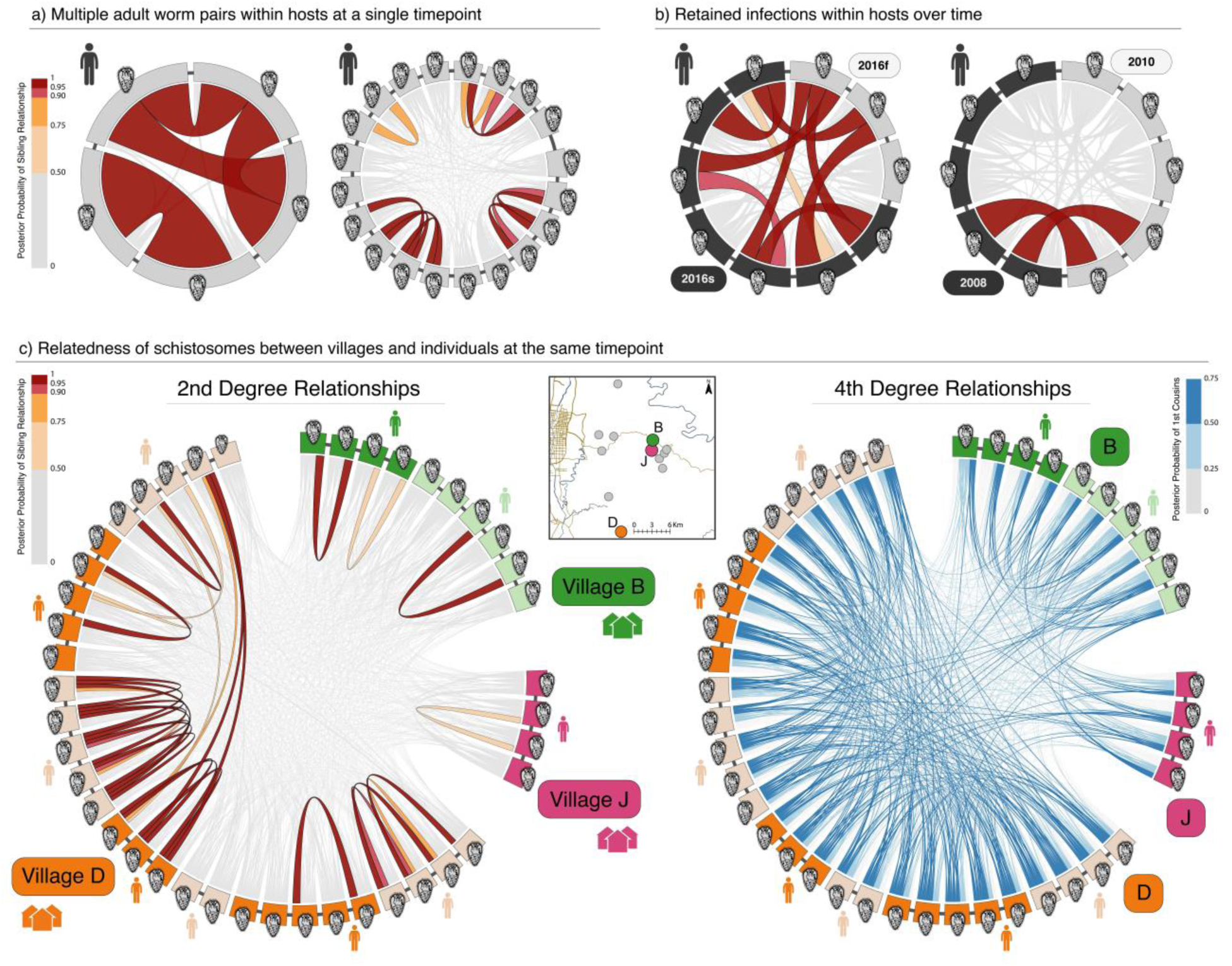
Relatedness of miracidia within and between hosts. Examples highlight relatedness structures indicating multiple infections, retained infections, evidence for clones, and inbreeding within villages. Hosts are indicated with human figures, with different miracidia collected from a single host connected by thin dark grey lines. Ribbons between miracidia show posterior probabilities of degree of relatedness through color (scale to side) and by ribbon width. a) Two examples of multiple sibling clusters (2 and 4) within hosts are shown. In the second example, 8 miracidia are not in sibling clusters (all connections are in light grey), indicating a high multiplicity of infection sources (a minimum of 12 mating pairs) in this host. b) Two examples of likely retained infections over time are inferred from the sibling-level miracidia sampled from the same host at different timepoints separated by five months and two years. c) Miracidia from multiple hosts living in villages B, J, and D and sampled at the same timepoint are shown with gaps between different villages. In the left graph, sibling-level relatedness is shown, and a case of sibling-level relatedness between miracidia across two individuals indicates clonal parents. In the right graph, cousin-level relatedness emphasizes that strongly supported first-cousin relationships are common among miracidia within villages and sparse between villages.

The most direct human-to-human infection pathway, involving only a snail intermediate host, would yield avuncular relationships between miracidia from each host. Thus, evidence for an avuncular relationship would provide the clearest evidence to date of a human reservoir. We are unable to differentiate between the types of 2^nd^ degree relations (double first cousins, half siblings, or avuncular), but all three of these relationships provide evidence of a small schistosome population size and high inbreeding. Interpreting broad patterns of 2^nd^ degree relationships is complicated by the lifecycle of schistosomes. The human-infective cercaria stage of schistosomes that originates from snails is clonal, and genetically-identical cercariae may produce multiple infections in one or more human hosts (*19*). Although rare, our observation of sibling-level relatives across human hosts demonstrates multiple clonal infections have occurred (Fig 3c), and the offspring of clonal infections could explain some 2^nd^ degree relatives observed between human hosts.

While the existence of clones can theoretically produce false inferences of sibling relationships (and thus retained infections) within individual hosts across time points, we discount this partly due to the ∼6-month lifespan of infected *Oncomelania* snails (*20*) and the months required to develop worms from cercariae and form mating pairs within a human host. Time-separated clonal double infections would require the environmental condition that the individual snails produce clonal cercariae from the time of the initial infection (followed by development, mate pairing, detection, and treatment- a minimum of 40 days (*21*)) until the time of the second infection. In contrast, the retention of infections over time, due either to non-compliance with treatment or treatment failure, is a more obvious explanation and supported by prior evidence (*22*). Furthermore, cross-host siblings indicating clonal pairs are rare (Fig 3c). Thus, we generally expect that clonality has had little impact on our within-host inferences of retained infection. Instead, the detection of schistosome clones between human hosts suggests that schistosomiasis control efforts have been highly effective in reducing local snail populations required for producing cercariae. These findings illustrate how control measures targeting the reduction of snail populations have shaped schistosome reproductive and transmission patterns that have in turn impacted genetic structure and relatedness.

The work presented here exemplifies how population genomic studies can illuminate factors underlying transmission and persistence of macroparasites and provide strategic and precise advice to direct control efforts. We find that there are high levels of schistosome inbreeding within villages, that local sources of infection are consistent through time, and that there is occasional persistent infection in individual humans despite treatment referral. These findings indicate that the persistence of schistosomiasis in residual transmission hotspots is likely due to local transmission and reinfection, with at least some contribution from humans. Based on our findings, end-game control measures should focus on confirmation of schistosome elimination from infected human hosts and complete extirpation of local infection reservoirs.

## Acknowledgments

We thank the members of the Sichuan Centers of Disease Control and the local county anti-schistosomiasis stations for their assistance in collecting parasite samples and related field data.

## Author contributions

Conceptualization- TAC, EJC, DDP; Ideas- TAC, EJC, DDP; Methodology-JAS, TAC, EJC, DDP; Software- JAS, DDP; Validation- LET; Formal analysis- JAS, LET, ZLN, DRS, BWP, TAC, DDP; Investigation- JAS, LET, NRH, TAC, DDP; Resources- BZ, YL, EJC, TAC, DDP; Data Curation- JAS, LET, TAC, DDP; Writing – original draft preparation- JAS, EJC, DDP; Writing – review and editing- JAS, LET, ZLN, DRS, BWP, YL, BZ, TAC, EJC, DDP; Visualization- JAS, LET, ZLN, DRS, BWP, TAC, DDP; Supervision- TAC, EJC, DDP; Project administration- TAC, EJC, DDP; Funding acquisition- YL, BZ, EJC, TAC, DDP.

## Data and materials availability

Sequences generated during this work have been deposited in the NCBI Sequence Read Archive under BioProject PRJN xxxxxxx The vcf file used in analysis, as well as custom scripts are available at www.evolutionarygenomics.com/ProgramsData/SchistoGenomics/

## Supplementary Materials

Materials and Methods

Figures S1-S3

Table S1

References (*23-35*)

