## Supplementary Material for "Population genomic analyses of schistosome parasites highlight critical challenges facing end-game elimination efforts"

**This PDF file includes:**

Materials and Methods
Supplementary Text
Figs S1 to S3
Table S1

### Materials and Methods

Miracidia collection and sample selection

Miracidia, the progeny of adult schistosomes, were collected from 12 villages in Sichuan, China (see Figure 1A). Infection surveys took place in 2007, 2008, 2010, and in both the summer and fall of 2016. During each survey village residents submitted fecal samples for three consecutive days and each sample was tested for *S. japonicum* infection using the miracidium hatching test as described elsewhere (*13*). Individual miracidia were collected from the top of the hatching test flask, rinsed three times in autoclaved de-ionized water and transferred to Whatman FTA indicator cards using a hematocrit tube or Pasteur pipette drawn to a narrow bore with a flame.

A subset of collected samples were selected for inclusion in the study. This subsampling was designed to include 10-15 miracidia from every village and across multiple timepoints. When possible, we tried to include multiple samples from the same human host.

The research involving human subjects was approved by the Sichuan Institutional Review Board, the University of California, Berkeley, Committee for the Protection of Human Subjects, and the Colorado Multiple Institutional Review Board. Participants provided written, informed consent. Anyone testing positive for *Schistosoma japonicum* was informed of their infection status and referred to the local anti-schistosomiasis control station for treatment.

DNA library preparation and sequencing

DNA library preparation followed methodology described in (*14*). Briefly, discs containing individual miracidia were excised from Whatman FTA cards using a 2mm card punch (Whatman WB100029) and DNA from the disc was whole-genome-amplified by isothermal genome amplification, termed "multiple displacement amplification" (MDA), using GenomiPhi v3 (GE Healthcare Biosciences 25660124) amplification tubes with modifications described in (*14*). Amplified DNA was digested for >8 hrs with *PstI-HF* and *Sau3AI* at 37°C followed by a 65°C heat deactivation step. Following solid phase reverse immobilization (SPRI) DNA extraction, custom adaptors containing an 8-bp unique molecular identifier (UMI) and sequences corresponding to the single-stranded DNA sticky ends generated by digestion and a 6-bp barcode were ligated to digested fragments. Adaptors ligating to *PstI-HF* cuts also contained 6 bp barcodes. Adaptor sequences used can be found in Table S1. Following ligation, sets of 6−8 samples were pooled such that no barcodes were used twice within the same pool, and underwent size selection for fragments sizes either between 300 and 600 bp (including adaptor sizes) or 390 and 690 bp (including adaptor sizes) using a PippinPrep with a 1.5% Agarose gel. Following size selection, samples underwent 15 cycles of PCR amplification. Primers used in amplification also contained index sequences and sequences used for illumina-based sequencing cluster formation. Sample pools were then combined in equimolar ratios such that no index sequence was used more than once within each pool. Samples were sequenced on an Illumina HiSeq using v4 chemistry. In total 272 samples were sequenced: 124 with 125 bp single end reads, and 148 of with 150 bp paired end reads. In total 1,799,089,548 reads were generated.

Fastq processing and variant identification

PCR clones were filtered from the reads with the clone filter tool in s*tacks* (*23*) using the UMIs contained in each barcode, with an average of 5.61% of reads filtered out of each library as clones. Sequences were then quality filtered and divided by barcode using the *process-radtags* tool in *stacks* (*23*) with restriction enzymes and barcodes supplied as arguments as well as the options q, r, and D, which rescue single base mutations in restriction sites and barcodes, remove reads without restriction sites in the correct location and reads with unclear barcodes, and filter reads with low quality using a sliding window read quality score. An average of 31.37% of reads were filtered from each library due to clonality, ambiguous barcodes/restriction site, or quality, though one library containing barcoded DNA from eight miracidia contained an abnormally high number of reads missing restriction sites in the correct place. This library was retained, with the filters above applied. Excepting this library, an average of 26.96% of total reads were filtered from all reads. Reads were mapped to the *S. japonicum* reference genome (downloaded from schistodb.net (*24, 25*) using *bwa mem* (*26*). Variants were called from .bam files using *Haplotype Caller* in the Genome Analysis Toolkit (*27-29*) with gvcf mode and GenotypeGVCFs. Over 4 million variant sites were found, but most of these sites were sequenced in just one or very few miracidia. Demultiplexed fastq files, as well as bam and bam index files, are available through the NCBI SRA database under BioProject PRJN xxxxxxx.

Defining sets of ddRADseq loci

Although most ddRADseq reads mapped reliably to expected ddRADseq loci (*14*), loci from off-target reads may add noise to subsequent analyses. We therefore undertook to identify a set of loci that were reliably recovered at sufficient depth across the majority of samples in order to retain only those variants that map to ddRADseq loci. Using a custom *perl* script (*cutgenome.pl*; EvolutionaryGenomics.com/ProgramsData/SchistoGenomics), we identified the expected mapping locations of ddRADseq reads in the *S. japonicum* reference genome (*25*), with each expected individual ddRADseq locus having two different locations: one for the forward read and one for the reverse read, if applicable.

To identify the subset of these expected loci that could be reliably recovered, we first eliminated miracidia that had fewer than 500K reads post-filtering or less than 20K reads that map to the reference genome with a mapq ≥ 20. This step mitigates effects caused by poor initial amplification, or errors in library preparation or sequencing in determining which ddRADseq loci can be reliably recovered. We obtained the sequencing depth of each expected ddRADseq locus in each of the 156 remaining ‘high-depth’ miracidia using *bedtools* (*30*) intersect with the -c and -f 0.5 arguments (-c to record the coverage of the expected read locus and -f 0.5 to ensure that mapped loci overlapped by at least 50% of an expected read locus before incrementing the depth count). Depending on the sequencing protocol, we applied different criteria to determine the depth of each ddRADseq locus for miracidia that were sequenced with single and paired-end reads. For miracidia that were sequenced with single end sequencing, the cumulative depth of each of the fragment’s possible reads was used; for miracidia sequenced with paired end sequencing, the mean depth of the two read loci was used for the fragment’s depth. 9,637 expected ddRADseq loci of any size were sequenced at ≥20x depth in ≥75% of ‘high-depth’ miracidia (see Fig S1). To further restrict variants to the most stringent loci, analyses reported here used only variants from the 6,990 expected ddRADseq loci that were close to the target size selection range (170bp−500bp). The bed file containing this data set can be downloaded from EvolutionaryGenomics.com/ProgramsData/SchistoGenomics.

Variation set creation

Variants then underwent a number of filters as follows: invariant sites were removed, sites with more than two alleles or that contained an indel were removed, and variants that were not within an expected ddRADseq locus were removed. After applying these filters, 72,797 variants remained. To create our final variant set, we re-coded any sites sequenced at <10x coverage to missing data, recoded individual genotypes with GQ <20 as missing, and removed sites that were missing more than 75% of genotype calls. This variant set contained 33,901 variants in 200 miracidia. The vcf file containing this data set is available at EvolutionaryGenomics.com/ProgramsData/SchistoGenomics.

Population analyses

The parametric tests for population structure we performed require that the provided loci be in linkage equilibrium, however missing genetic distances between neighboring sites and the currently highly-fragmented reference genome makes linkage pruning difficult. This problem is compounded in our dataset because a large proportion of the miracidia were suspected to be highly related which could inflate linkage estimates between sites. Here, we outline the steps we performed to obtain a reasonably unlinked set of variants though we note that this pruning does not guarantee that all sites used are in linkage equilibrium.

We first identified miracidia that are expected to be closely related by identifying clusters of miracidia that share a proportion of rare alleles greater than 0.45 (see 'Identification of family clusters and relatedness estimates' below) between each pair of miracidia, and removed all but one miracidium from each cluster. A total of 83 miracidia remained following this step. We then pruned linked variants in this putative unrelated set using *plink*'s –indep-pairwise command (v1.90b4.6; *31*) with arguments 1000 100 0.1, which greedily prunes variants with $r^2 > 0.1$ from overlapping windows consisting of 1,000 variants. Using a low $r^2$ threshold with a window size large enough to encompass whole ddRAD fragments reduces the set of variants on each ddRAD fragment to the subset of variants that are in relative linkage equilibrium, with the goal that such strict filters might at least partially mitigate the undesired effects of the inability to test for linkage between variants residing on different contigs of the reference genome. Linkage pruning in this way reduced the number of variants in the putatively unrelated set to 6,642.

We used ADMIXTURE (*16*) and these putatively unlinked variants with all 200 miracidia to determine the proportion of each miracidium's genome that can be attributed to one of $k$ different populations. We tested values for $k$ ranging from 2 to 10 with ten replicates for each $k$ and default cross-validation to determine the $k$ with the lowest cross validation error.

Principal component analysis (PCA), as implemented in *R*'s SNPrelate package (*32*), was applied to the full variant set to assess how genotype differences between miracidia contribute to region-wide variability between samples and villages.

Using all variants, we calculated pairwise genetic distances between miracidia through the distance-based *bitwise.dist* function implemented in *R*'s adegenet package (*33, 34*) and used distances to construct a neighbor-joining tree using the *R* package 'ape' (*35*). We then visualized relationships based on both time and village to further illustrate structuring across samples.

Identification of family clusters and relatedness estimates

To identify highly related samples in the absence of reliable allele frequency estimates, pairwise comparison of rare allele sharing (MAF ≤ 0.1, MAF based on frequency of allele in whole data set) (RAS) was calculated between all pairs of samples using only rare variants and a custom perl script (findSibClusters.pl; EvolutionaryGenomics.com/ProgramsData/SchistoGenomics) following

$$P_{ij} = \frac{1}{L}\sum_{k=1}^{L} x_{ijk}$$

where

$$x_{ijk} = \begin{cases} 1\ if\ \ i\ and\ j\ have\ the\ same\ genotype\ at\ locus\ k \\ 0.5\ if\ \ i\ and\ j\ share\ one\ allele\ at\ locus\ k \\ 0\ if\ \ i\ and\ j\ share\ no\ alleles\ at\ locus\ k \end{cases}$$

and $P_{ij}$ is the proportion of shared alleles between individuals $i$ and $j$, $L$ is the number of loci tested, and $k$ is a locus for which both individuals $i$ and $j$ have non-missing genotype calls and individual $i$ has a rare variant. To avoid overestimating relationships because of linked variants, we use the mean proportion of rare alleles shared generated from 30 replicates of randomly sampling 2,000 loci with replacement for each pairwise comparison. We identified clusters of highly related miracidia such that each miracidium in a cluster shared ≥ 0.45 of its rare alleles with at least one other miracidium in the cluster. Removal of all-but-one miracidia from each putative sibling cluster (117 individuals) resulted in a data set of 83 miracidia. The sibling-pruned vcf file is available at EvolutionaryGenomics.com/ProgramsData/SchistoGenomics.

<u>Calculating posterior probabilities across degrees of relatedness</u>

To generate posterior probability distributions for each degree of relatedness, we first estimated mean levels of unrelated allele sharing, $\hat{m}_{unrelated}$, as the average pairwise rare allele sharing between individuals from the most geographically distant villages (n=35.6km). To estimate allele sharing among clusters of miracidia likely to be siblings (1st degree relatives), we included pairs from clusters of 3 or more miracidia from the same host all with rare allele sharing proportion >= 0.30 (45 miracidia in 13 clusters). There were an additional 8 pairs of miracidia within the same host that are likely siblings but not part of a big enough cluster. The estimated mean, $\hat{m}_{sibs}$ and variance, $\hat{s}_{sibs}$, of allele sharing were calculated from eligible pairs (n = 60). For intermediate degrees of relatedness, means were estimated by successively halving the distance from sibs to unrelated, and variances were estimated by successively halving the sibling variance for each further degree of relatedness (e.g., $\hat{m}_{2^\circ} = \frac{\left(\hat{m}_{unrelated} + \hat{m}_{sibs}\right)}{2}$ and $\hat{s}_{2^\circ} = \frac{\hat{s}_{sibs}}{2}$). Posterior probabilities were calculated roughly assuming even prior probabilities for each categorical degree of relatedness from siblings to 5th degree relatives and unrelated, and assuming that allele sharing probabilities for each degree of relatedness were distributed normally, i.e, $\sim N\left(\hat{m}_{degree}, \hat{s}_{degree}\right)$.

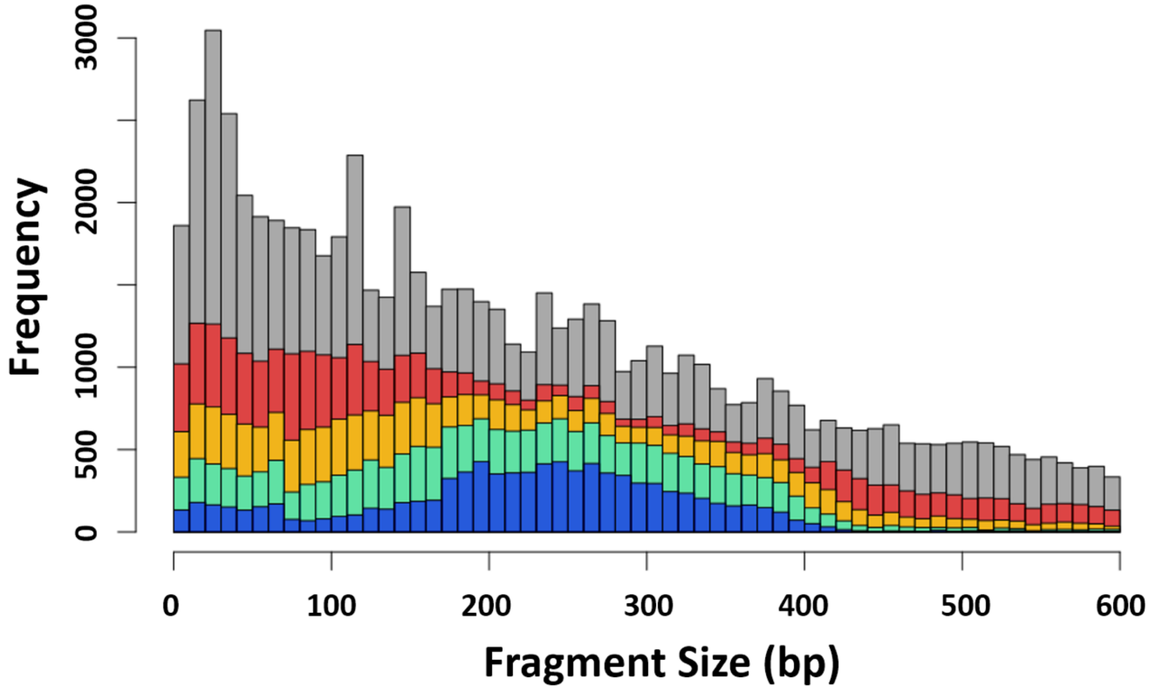


**Fig. S1.**

Size distribution of expected ddRADseq loci. Size distribution of expected ddRADseq loci sequenced in at least 75% of 'high-depth' miracidia (see text) at different depths. Gray = all ddRAD fragments, red = 2x coverage, orange = 5x coverage, green = 10x coverage, blue = 20x coverage.

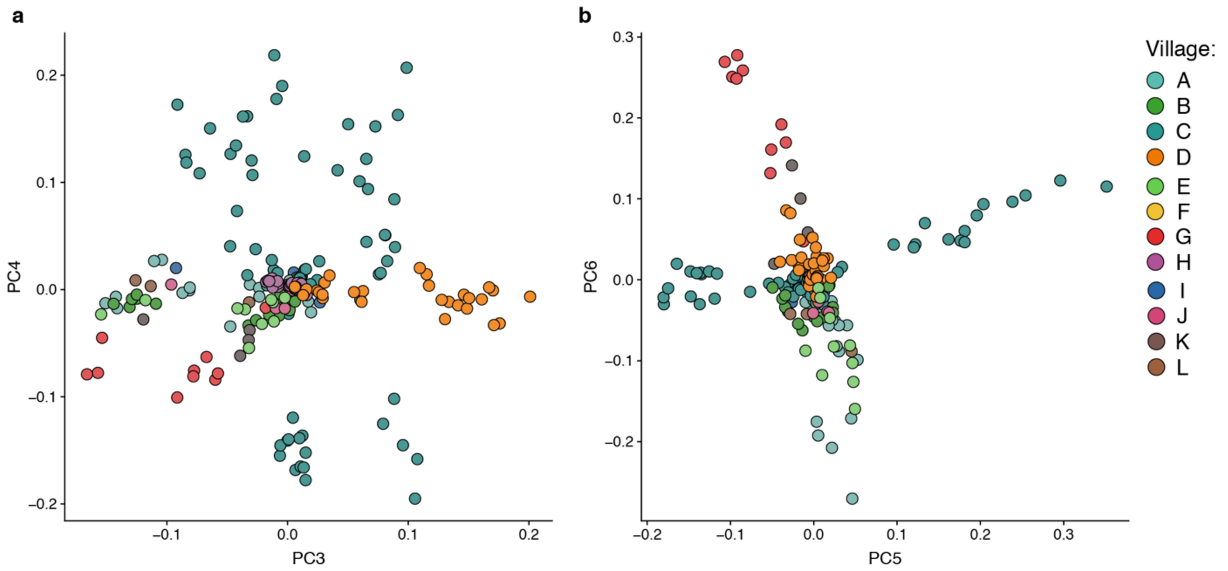


**Fig. S2.**

Principal components analysis of 200 miracidia from 12 villages. Each point corresponds to an individual miracidium, with color of the points indicating the village from which the miracidium was collected. PC's 3 and 4 are shown in the figure on the left and PC's 5 and 6 are shown in the figure on the right. Village colors correspond to those shown in Fig 1a.

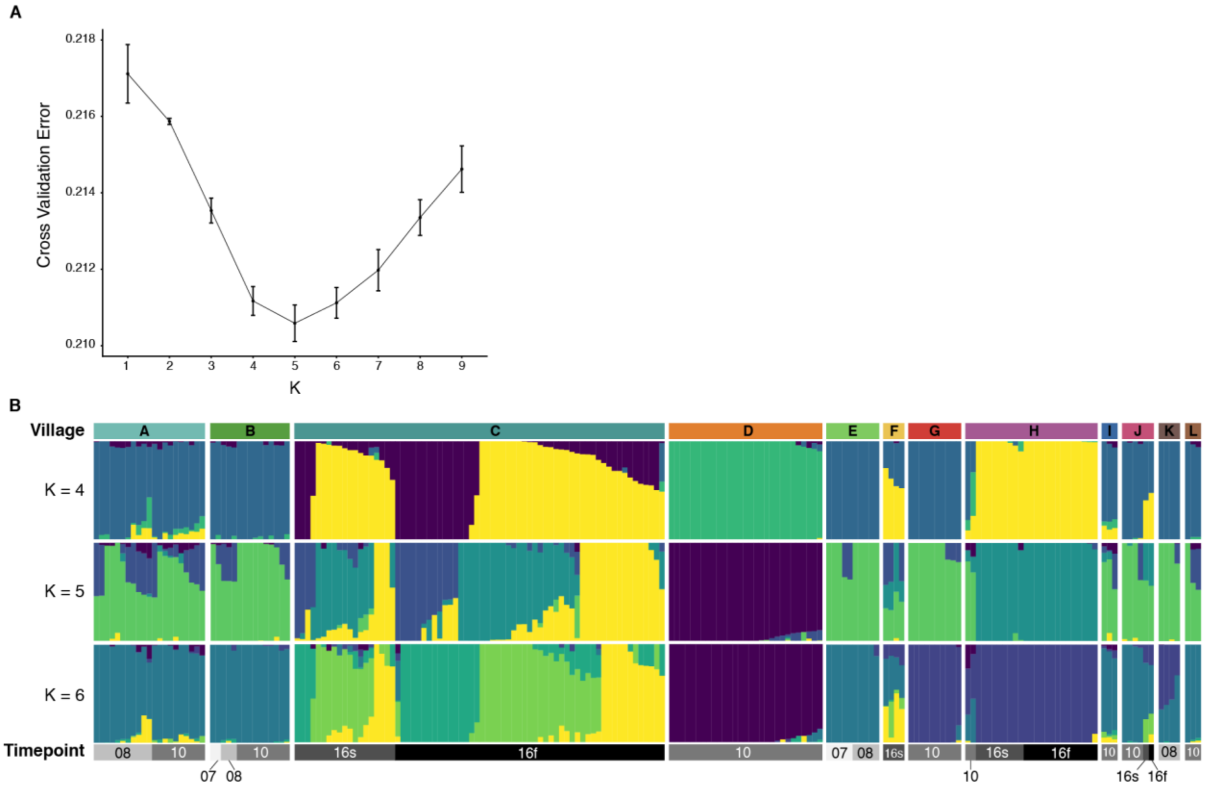


**Fig. S3.**

Supplemental ADMIXTURE plots. As in Fig 1e, ADMIXTURE plot showing genetic clusters grouped by village and sampling timepoint. A) Cross-validation error for different values *k* based on 10 runs at each *k*. Error bars represent standard error of the mean for each cross-validation value at the specified *k*. B) graphical results for *k*=4, 5, and 6 for comparison. *k*=4 was shown in figure 1.

Table S1. ddRADseq adaptor and primer oligonucleotides sequences.

| Type | ID | Sequence (5' to 3') |
|---|---|---|
| Adaptor | Barcode1 | ACACTCTTTCCCTACACGACGCTCTTCCGATCTNNNNNNNNACTAGGTGCA<br>CCTAGTNNNNNNNNAGATCGGAAGAGCGTCGTGTAGGGAAAGAGTGT |
| Adaptor | Barcode2 | ACACTCTTTCCCTACACGACGCTCTTCCGATCTNNNNNNNNAGCATTTGCA<br>AATGCTNNNNNNNNAGATCGGAAGAGCGTCGTGTAGGGAAAGAGTGT |
| Adaptor | Barcode3 | ACACTCTTTCCCTACACGACGCTCTTCCGATCTNNNNNNNNAGTAAGTGCA<br>CTTACTNNNNNNNNAGATCGGAAGAGCGTCGTGTAGGGAAAGAGTGT |
| Adaptor | Barcode4 | ACACTCTTTCCCTACACGACGCTCTTCCGATCTNNNNNNNNATAACCTGCA<br>GGTTATNNNNNNNNAGATCGGAAGAGCGTCGTGTAGGGAAAGAGTGT |
| Adaptor | Barcode5 | ACACTCTTTCCCTACACGACGCTCTTCCGATCTNNNNNNNNATGTCCTGCA<br>GGACATNNNNNNNNAGATCGGAAGAGCGTCGTGTAGGGAAAGAGTGT |
| Adaptor | Barcode6 | ACACTCTTTCCCTACACGACGCTCTTCCGATCTNNNNNNNNCAGAGTTGCA<br>ACTCTGNNNNNNNNAGATCGGAAGAGCGTCGTGTAGGGAAAGAGTGT |
| Adaptor | Barcode7 | ACACTCTTTCCCTACACGACGCTCTTCCGATCTNNNNNNNNCATCTCTGCA<br>GAGATGNNNNNNNNAGATCGGAAGAGCGTCGTGTAGGGAAAGAGTGT |
| Adaptor | Barcode8 | ACACTCTTTCCCTACACGACGCTCTTCCGATCTNNNNNNNNAACGGTTGCA<br>ACCGTTNNNNNNNNAGATCGGAAGAGCGTCGTGTAGGGAAAGAGTGT |
| Adaptor | Barcode9 | ACACTCTTTCCCTACACGACGCTCTTCCGATCTNNNNNNNNCCCATATGCA<br>TATGGGNNNNNNNNAGATCGGAAGAGCGTCGTGTAGGGAAAGAGTGT |
| Adaptor | Barcode10 | ACACTCTTTCCCTACACGACGCTCTTCCGATCTNNNNNNNNCGAAACTGCA<br>GTTTCGNNNNNNNNAGATCGGAAGAGCGTCGTGTAGGGAAAGAGTGT |
| Adaptor | Barcode11 | ACACTCTTTCCCTACACGACGCTCTTCCGATCTNNNNNNNNCGAATGTGCA<br>CATTCGNNNNNNNNAGATCGGAAGAGCGTCGTGTAGGGAAAGAGTGT |
| Adaptor | Barcode12 | ACACTCTTTCCCTACACGACGCTCTTCCGATCTNNNNNNNNGACCAATGCA<br>TTGGTCNNNNNNNNAGATCGGAAGAGCGTCGTGTAGGGAAAGAGTGT |
| Adaptor | Barcode13 | ACACTCTTTCCCTACACGACGCTCTTCCGATCTNNNNNNNNGACGTTTGCA<br>AACGTCNNNNNNNNAGATCGGAAGAGCGTCGTGTAGGGAAAGAGTGT |
| Adaptor | Barcode14 | ACACTCTTTCCCTACACGACGCTCTTCCGATCTNNNNNNNNGCAGAATGCA<br>TTCTGCNNNNNNNNAGATCGGAAGAGCGTCGTGTAGGGAAAGAGTGT |
| Adaptor | Barcode15 | ACACTCTTTCCCTACACGACGCTCTTCCGATCTNNNNNNNNGGGATATGCA<br>TATCCCNNNNNNNNAGATCGGAAGAGCGTCGTGTAGGGAAAGAGTGT |
| Adaptor | Barcode16 | ACACTCTTTCCCTACACGACGCTCTTCCGATCTNNNNNNNNGTCTATTGCA<br>ATAGACNNNNNNNNAGATCGGAAGAGCGTCGTGTAGGGAAAGAGTGT |
| Adaptor | Barcode17 | ACACTCTTTCCCTACACGACGCTCTTCCGATCTNNNNNNNNTAAGACTGCA<br>GTCTTANNNNNNNNAGATCGGAAGAGCGTCGTGTAGGGAAAGAGTGT |
| Adaptor | Barcode18 | ACACTCTTTCCCTACACGACGCTCTTCCGATCTNNNNNNNNTCAATCTGCA<br>GATTGANNNNNNNNAGATCGGAAGAGCGTCGTGTAGGGAAAGAGTGT |

| | | |
|---|---|---|
| Adaptor | Barcode19 | ACACTCTTTCCCTACACGACGCTCTTCCGATCTNNNNNNNNTCTGCTTGCA<br>AGCAGANNNNNNNNAGATCGGAAGAGCGTCGTGTAGGGAAAGAGTGT |
| Adaptor | Barcode20 | ACACTCTTTCCCTACACGACGCTCTTCCGATCTNNNNNNNNTCTTAGTGCA<br>CTAAGANNNNNNNNAGATCGGAAGAGCGTCGTGTAGGGAAAGAGTGT |
| Adaptor | Barcode21 | ACACTCTTTCCCTACACGACGCTCTTCCGATCTNNNNNNNNTGCTCATGC*A<br>TGAGCANNNNNNNNAGATCGGAAGAGCGTCGTGTAGGGAAAGAGTGT |
| Adaptor | Barcode22 | ACACTCTTTCCCTACACGACGCTCTTCCGATCTNNNNNNNNTGGGATTGCA<br>ATCCCANNNNNNNNAGATCGGAAGAGCGTCGTGTAGGGAAAGAGTGT |
| Adaptor | Barcode23 | ACACTCTTTCCCTACACGACGCTCTTCCGATCTNNNNNNNNTGTTGGTGCA<br>CCAACANNNNNNNNAGATCGGAAGAGCGTCGTGTAGGGAAAGAGTGT |
| Adaptor | Barcode24 | ACACTCTTTCCCTACACGACGCTCTTCCGATCTNNNNNNNNACCAAATGCA<br>TTTGGTNNNNNNNNAGATCGGAAGAGCGTCGTGTAGGGAAAGAGTGT |
| Adaptor | Universal | GTGACTGGAGTTCAGACGTGTGCTCTTCCGATCTNNNNNNNN<br>GATCNNNNNNNNAGATCGGAAGAGCGAGAACAA |
| Primer | Index1 | CAAGCAGAAGACGGCATACGAGATCGTGATGTGACTGGAGTTCAGACGTGTGC |
| Primer | Index2 | CAAGCAGAAGACGGCATACGAGATACATCGGTGACTGGAGTTCAGACGTGTGC |
| Primer | Index3 | CAAGCAGAAGACGGCATACGAGATGCCTAAGTGACTGGAGTTCAGACGTGTGC |
| Primer | Index4 | CAAGCAGAAGACGGCATACGAGATTGGTCAGTGACTGGAGTTCAGACGTGTGC |
| Primer | Index5 | CAAGCAGAAGACGGCATACGAGATCACTGTGTGACTGGAGTTCAGACGTGTGC |
| Primer | Index6 | CAAGCAGAAGACGGCATACGAGATATTGGCGTGACTGGAGTTCAGACGTGTGC |
| Primer | Index7 | CAAGCAGAAGACGGCATACGAGATGATCTGGTGACTGGAGTTCAGACGTGTGC |
| Primer | Index8 | CAAGCAGAAGACGGCATACGAGATTCAAGTGTGACTGGAGTTCAGACGTGTGC |
| Primer | Index9 | CAAGCAGAAGACGGCATACGAGATCTGATCGTGACTGGAGTTCAGACGTGTGC |
| Primer | Index10 | CAAGCAGAAGACGGCATACGAGATAAGCTAGTGACTGGAGTTCAGACGTGTGC |
| Primer | Index11 | CAAGCAGAAGACGGCATACGAGATGTAGCCGTGACTGGAGTTCAGACGTGTGC |
| Primer | Index12 | CAAGCAGAAGACGGCATACGAGATTACAAGGTGACTGGAGTTCAGACGTGTGC |
| Primer | Universal | AATGATACGGCGACCACCGAGATCTACACTCTTTCCCTACACGACG |
